# Inhibition of hexokinase 2 undermines cartilage health and accelerates osteoarthritis

**DOI:** 10.64898/2026.05.26.728030

**Authors:** Haokun Xu, Xinyi Zhang, Yijie Fu, Genming Liu, Sirui Yuan, Daizhao Deng, Kai Li, Tinghui Xiao, Ying Lin, Ruijun Lai, Song Xu, Xiaochun Bai, Yue Zhang

## Abstract

**Objective:** Enhanced glycolysis is a metabolic hallmark of chondrocytes in osteoarthritis (OA); however, the roles of the glycolytic rate-limiting enzyme hexokinase 2 (HK2) in cartilage remain poorly understood.

**Methods:** Pharmacological approach (3-bromopyruvate (3-BrPA) treatment) and mice model involving HK2 knockout in Col2a1-expressing chondrocytes are utilized to access the impact of HK2 blockage on cartilage *ex vivo* and *in vivo*. The *in vivo* effects of HK2 inhibition on OA progression were evaluated using a destabilization of the medial meniscus (DMM)-induced OA mouse model, through both intra-articular 3-BrPA administration and chondrocyte HK2 deletion. Additionally, we analyzed published single-cell RNA sequencing (scRNA-seq) datasets from human articular cartilage and integrated these with bulk RNA-seq data from HK2-deficient chondrocytes to characterize HK2 expression features across conditions.

**Results:** Both pharmacological inhibition and genetic deletion of HK2 impair cartilage formation *ex vivo*. Bulk RNA-seq analysis and *ex vivo* studies demonstrated a promoted ossification-like process due to HK2 ablation in chondrocytes. Through pseudotime analysis of published single-cell RNA sequencing (scRNA-seq) datasets from human articular cartilages, we further identified that HK2 is differentially expressed across conditions, with a feature of a relatively high expression level at terminal stages of chondrocyte differentiation in the context of OA. We next confirmed HK2 deficiency in chondrocytes significantly exacerbated OA progression but having no impact on skeletal development in mice.

**Conclusions:** HK2 plays a critical role in maintaining cartilage health, likely through the regulation of calcification, thereby highlighting the potential risks associated with targeting glycolytic enzymes as a therapeutic strategy for OA.

## Introduction

Cartilage functional loss is the main characteristics for the progressive development in osteoarthritis (OA) [1, 2]. Emerging evidence suggests that metabolic reprogramming may provide a potential therapeutic target for impeding cartilage dysfunction in OA management [3, 4], however, the roles of metabolic enzymes involve supporting cartilage formation and function remains poorly understood.

In healthy joints, chondrocytes maintain metabolic homeostasis to support normal function. Under OA conditions, alterations in the joint microenvironment lead changes in chondrocyte carbon metabolism [5, 6]. Growing evidence indicates a metabolic shift toward glycolysis in OA chondrocytes, which may promote inflammation and contribute to disease progression [4, 7]. Enhancing mitochondrial respiration has shown promise in restoring cartilage and joint function in OA [3, 8]. Moreover, inhibiting glycolysis appears to attenuate the progression of rheumatoid arthritis (RA) [9, 10], a chronic autoimmune disease that primarily affects joints [11]. These findings might prompt a great deal of therapeutic attempts to target glycolysis for OA management.

However, whether glycolytic enzymes involved in chondrocyte homeostasis contributes to OA remains controversial [12, 13]. More importantly, several glycolytic enzymes, including glyceraldehyde-3-phosphate dehydrogenase [14, 15], hexokinase [16, 17] and pyruvate kinase [18], have been reported to localize to mitochondrial membranes or translocate into the mitochondrial matrix under certain conditions, where they participate in regulating carbon flux. HK2, the rate-limiting enzyme in the first step of glycolysis, catalyzes the phosphorylation of glucose to produce glucose-6-phosphate [19]. While HK2-mediated metabolic pathways are known to regulate self-renewal and various cellular processes under both physiological and pathological conditions, such as cancer development [20, 21], hematopoiesis [22], and cardiac hypertrophy [23], its role in skeletal system homeostasis remains poorly understood.

In this study, we sought to elucidate the role of HK2 in chondrocytes and OA. Using publicly available scRNA-seq datasets, bulk RNA-seq analyses of HK2-deficeint chondrocytes along with *ex vivo* and *in vivo* experiments, we demonstrate that HK2 may regulate cartilage homeostasis by mainly affecting calcification, thereby HK2 blockage would exacerbate OA. Our findings caution a potential outcome on mitochondrial dysfunction caused by targeting glycolytic enzymes in OA management. However, the underling regulatory mechanism needs to be further elucidated.

## Materials and methods

### Mice and induction of OA

All mice were housed under specific pathogen-free conditions (22 °C, 60% humidity) with a 12-hour light/dark cycle, in accordance with the guidelines of the Southern Medical University Laboratory Animal Care. The Col2a1-cre transgenic mice line was a generous gift from Prof. Xiao Yang (Academy of Military Medical Sciences, Beijing, China). *HK2*-floxed mice were purchased from GemPharmatech (GemPharmatech Co., Ltd; catalog no. T009288). Col2-Cre^+/−^; *HK2*^flox/flox^ offspring, resulting in deletion of HK2 in chondrocytes. All mice were on a C57BL/6J background and bred under conditions with free access to food and water.

To eliminate the influence of estrogen on OA development [24], only male mice were used. OA was induced via destabilization of the medial meniscus (DMM) in 10-week-old mice. Mice were anesthetized with tribromoethanol, and sterile surgery was performed on the right knee. Following a medial parapatellar arthrotomy, the anterior fat pad was retracted to expose the medial meniscotibial ligament, which was then transected. The joint was irrigated with sterile saline, and the incision was closed. In sham-operation controls, the joint capsule was opened, but the ligament remained intact. Mechanical hypersensitivity to footpad stimulation was assessed weekly after surgery. At six weeks post-operation, joint tissues were collected for analysis. Cartilage degradation was assessed by Safranin O staining and scored using the Osteoarthritis Research Society International (OARSI) grading system [25].

### Isolation, culture and chondrocyte differentiation

Primary chondrocytes were isolated from the knee cartilage of 5-day-old mice following disinfection with 75% ethanol. Residual soft tissue was removed after digestion with trypsin for 1 hour. The tissue was then minced and digested with 0.2% collagenase II (C8000, Solarbio) for 6 hours. After centrifugation and filtration, cells were cultured in F12 medium (GIBCO, 21331020). Cells were maintained at 37 °C in a humidified incubator with 5% CO₂.

For *in vitro* differentiation, passage 2 chondrocytes were cultured in F12 medium supplemented with 10% fetal bovine serum (FBS), 10 nM vitamin C (Sigma, PHR008), 1 nM dexamethasone (Sigma, D4902), and 100 nM insulin-transferrin-selenium (ITS, GIBCO, 41400045). For experiments involving 3-bromopyruvate (3-BrPA; HY-19992, MedChemExpress), cells were treated with 5 μM 3-BrPA dissolved in DMSO. ITS-induced chondrogenesis was conducted for 7 days, with medium changes every other day.

### Immunoblotting

Total protein was extracted using ice-cold RIPA lysis buffer (KGP2100, KeyGEN) supplemented with a protease/phosphatase inhibitor cocktail and PMSF (Thermo Fisher Scientific). Protein concentration was determined using a BCA assay, and samples were separated by SDS-PAGE and transferred to PVDF membranes. The following primary antibodies were used (all at 1:1000 dilution): anti-COL2A1 (ab34712, Abcam), anti-MMP13 (A11148, ABclonal), anti-SOX9 (A19710, ABclonal), anti-COL1A1 (ab316222, Abcam), anti-RUNX2 (20700-1-AP, ProteinTech), anti-EGLN3 (82377-1-RR, ProteinTech), and anti-HK2 (bs-9455R, BIOSS). α-tubulin was used as an internal control. After incubation with HRP-conjugated secondary antibodies (1:1000 dilution), bands were visualized using enhanced chemiluminescence (ECL, P2300, New Cell & Molecular Biotech).

### Histological analysis and immunostaining

Knee joints were harvested, fixed in 4% paraformaldehyde, decalcified, embedded in paraffin, and sectioned at 4 μm. Sections were deparaffinized, rehydrated, and stained with Safranin O/Fast Green, hematoxylin and eosin (H&E), or TRAP. The severity of cartilage degeneration was assessed using the OARSI scoring system.

For immunohistochemistry (IHC) and immunofluorescence (IF), tissue sections were incubated with primary antibodies overnight at 4 °C, followed by appropriate secondary antibodies at room temperature.

### RNA extraction and qPCR

Total RNA was extracted from cultured cells or frozen tissue using TRIzol Reagent (9109 RNAiso Plus, Takara). The samples were lysed in 1 mL TRIzol and incubated for 10 minutes at room temperature. Chloroform (0.2 mL per 1 mL TRIzol) was added, and the mixture was shaken and incubated for 5 minutes. After centrifugation at 4 °C, the aqueous phase was collected and mixed with 0.5 mL isopropanol to precipitate RNA. Pellets were washed and dissolved in 50 μL nuclease-free water. RNA purity and concentration were measured using a NanoDrop spectrophotometer (Thermo Fisher Scientific).

Reverse transcription was performed using PrimeScript™ RT Master Mix (RR036Q, Takara). qPCR was conducted using ChamQ SYBR qPCR Master Mix (Q311-02, Vazyme). Each experiment included at least three biological replicates (independent RNA isolations) and two technical replicates per sample. Primer sequences are listed in Supplementary Table 1.

### Statistical analysis

All experiments were performed at least three times. Biological replicates refer to independent samples from different animals or cell isolations, while technical replicates refer to repeated measurements of the same sample. Data are presented as mean ± SEM. Statistical significance was assessed using an unpaired, two-tailed Student’s *t*-test for two-group comparisons and one-way ANOVA with Tukey’s post hoc test for multiple comparisons. The significance level of *p* < 0.05 was considered statistically significant. Graphs and statistical analyses were generated using GraphPad Prism 9.0 software.

## Results

### 3-BrPA and HK2 deletion both impar chondrocyte differentiation ex vivo

By *ex vivo* chondrogenic differentiation assays, quantitative PCR (q-PCR) analysis revealed that HK2 expression increased during the first three days post-induction, peaked at day 7, and then declined (Fig. 1A), suggesting a dynamic, probable stage-specific role for HK2 in chondrocyte differentiation. 3-Bromopyruvic acid (3-BrPA) [26] is a potent inhibitor of HK2, acting through direct suppression of its enzymatic activity. To investigate the role of HK2 in chondrocytes, we first performed *ex vivo* differentiation of primary chondrocytes. Toluidine blue staining revealed a significant reduction in the intensity and number of glycosaminoglycan (GAG)-positive cells following consecutive treatment with 5 mM 3-BrPA with chondrogenic induction (Fig. 1B). Q-PCR analysis demonstrated that 3-BrPA treatment downregulated expression levels of key molecular markers of chondrocyte differentiation across various stages, including *Sox9* (progenitor stage) (P = 0.0019), *Col2a1* (non-hypertrophic stage) (P <0.0001) and *Col10a1* (hypertrophic stage) (P = 0.0069) (Fig. 1C). Conversely, the expression of *Mmp13*, a marker of pre-ossification and matrix degradation, was significantly upregulated by 3-BrPA treatment (P <0.0001) (Fig. 1C). These findings were further supported by immunoblotting, which showed a marked decrease in Sox9 while an increase in Mmp13 protein levels following 3-BrPA treatment (Fig. 1D).

**Fig. 1.**
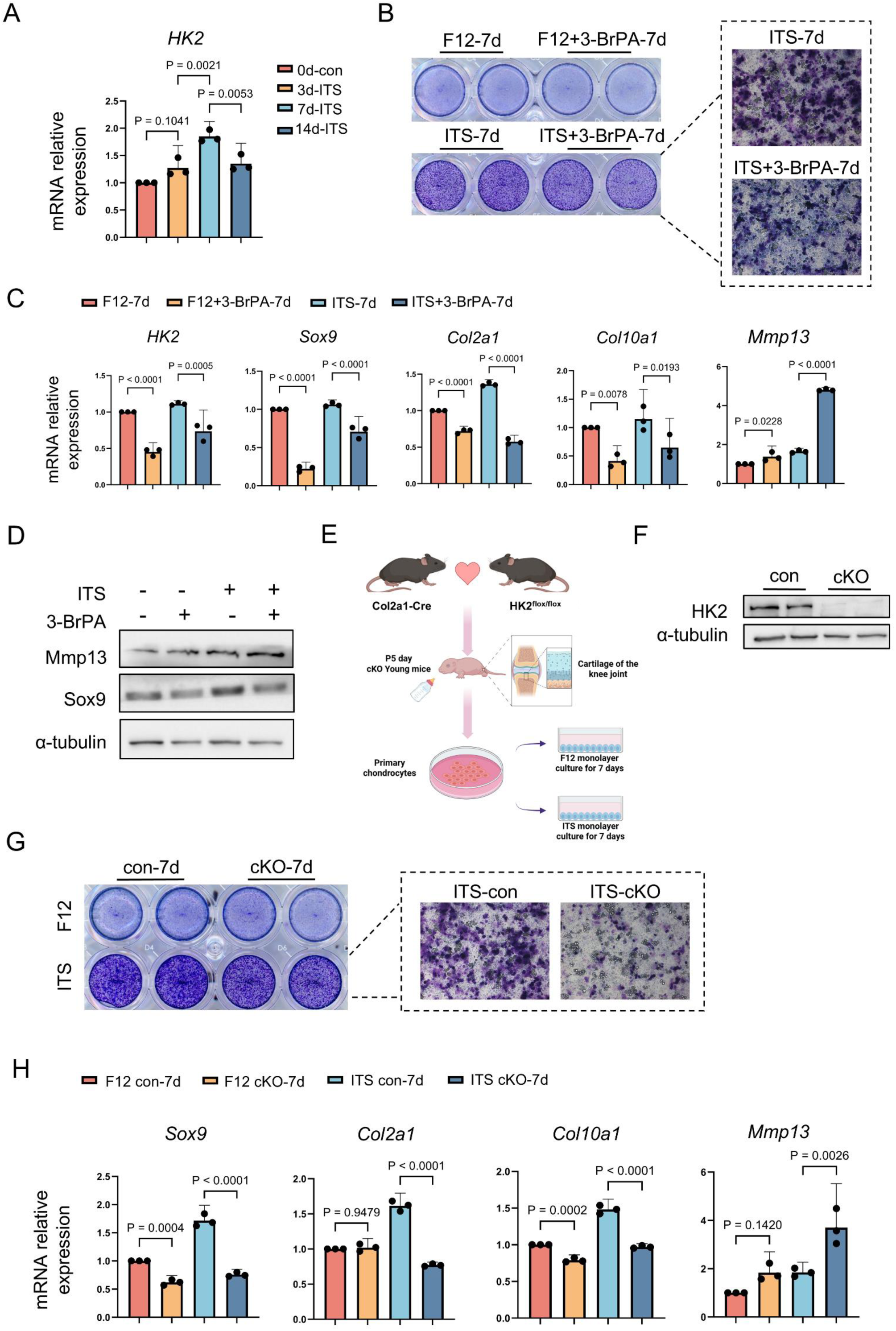
3-BrPA and HK2 deletion both impair chondrocyte differentiation *ex vivo*. (A) Relative mRNA expression levels of HK2 in chondrocytes treated with ITS for 0, 3, 7 and 14 days respectively. (B) Representative images of GAG-positive chondrocytes treated with or without 3-BrPA. Cells were differentiated towards the chondrogenic direction for 7 days by adding Insulin-Transferrin-Selenium (ITS). (C) Relative mRNA expression levels of *Sox9*, *Col2a1*, *Col10a1*, and *Mmp13* in chondrocytes treated with or without 3-BrPA under ITS induction. (D) Western blot analysis of Mmp13 and Sox9 protein levels in chondrocytes treated with or without 3-BrPA under ITS induction. (E) Schematic diagram of transgenic mouse model establishment and primary chondrocyte sampling. (F) Schematic of primary chondrocyte isolation from HK2 cKO and control mice, followed by ITS-induced differentiation. (G) Representative images of GAG-positive cells in HK2 cKO and control chondrocytes. Cells were differentiated towards the chondrogenic direction for 7 days by ITS induction. (H) Relative mRNA levels of *Sox9*, *Col2a1*, *Col10a1*, and *Mmp13* in HK2 cKO and control chondrocytes under ITS induction. All data are presented as mean ±SEM. *P < 0.05, **P < 0.01 indicates significant differences.

In order to confirm the particular effects of HK2 on chondrocyte differentiation, we generated a mice line with HK2-deficient chondrocytes (cKO). We conducted *ex vivo* culture and subsequent chondrogenic differentiation of primary chondrocytes isolated from HK2 cKO infant mice and their littermate controls (Fig. 1E, F). HK2-deficient chondrocytes exhibited impaired capacity in chondrogenic differentiation, as indicated by a reduced proportion of GAG-positive cells, even under chondrogenic induction (Fig. 1G), suggesting a resultant loss of proteoglycan due to HK2 ablation. Consistent with the effects observed following 3-BrPA treatment, HK2 deletion also resulted in decreased mRNA expression levels of *Sox9* (P <0.0001), *Col2a1* (P <0.0001), and *Col10a1* (P = 0.002), along with increased *Mmp13* (P = 0.0130) (Fig. 1H).

Given that Sox9 is a master transcription factor in driving chondrogenesis and Col2a1 expression, Col2a1 is the major structural component of the hyaline cartilage matrix and Col10a1 plays essential roles in endochondral ossification, the downregulated levels of genes related to matrix synthesis, and upregulated level of Mmp13 related to matrix degradation suggest that HK2 is essential for initiation of chondrogenic differentiation and the maintenance of the chondrocyte phenotype.

### HK2 ablation in chondrocytes suppresses energy generation and promotes calcification

Next, to investigate why HK2 ablation disrupts chondrocyte differentiation, we conducted *ex vivo* chondrogenic differentiation for 7 days, followed by transcriptomic analysis. Integrative analysis identified 1,194 differentially expressed genes (DEGs) (Supplemental Table 2), with 521 significantly upregulated and 673 downregulated (P < 0.05; Fig. 2A and Fig. S1A). The DEGs heatmap showed a marked downregulation of glycolytic genes, including *HK2* (Fig. 2B), as well as genes associated with mitochondrial function (Fig. 2C). Despite a significant enrichment of Kyoto Encyclopedia of Genes and Genomes (KEGG) in glycolysis (P < 0.0001) and oxidative phosphorylation (OXPHOS) (P < 0.0001) (Fig. 2D), gene set enrichment analysis (GSEA) only demonstrated significant suppression of OXPHOS (P = 0.0358) (Fig. 2E). Given that HK2 can locate on mitochondrial outer membrane involved in regulating mitochondrial homeostasis, these results suggest an advantage role of mitochondrial HK2 in chondrocytes.

**Fig. 2.**
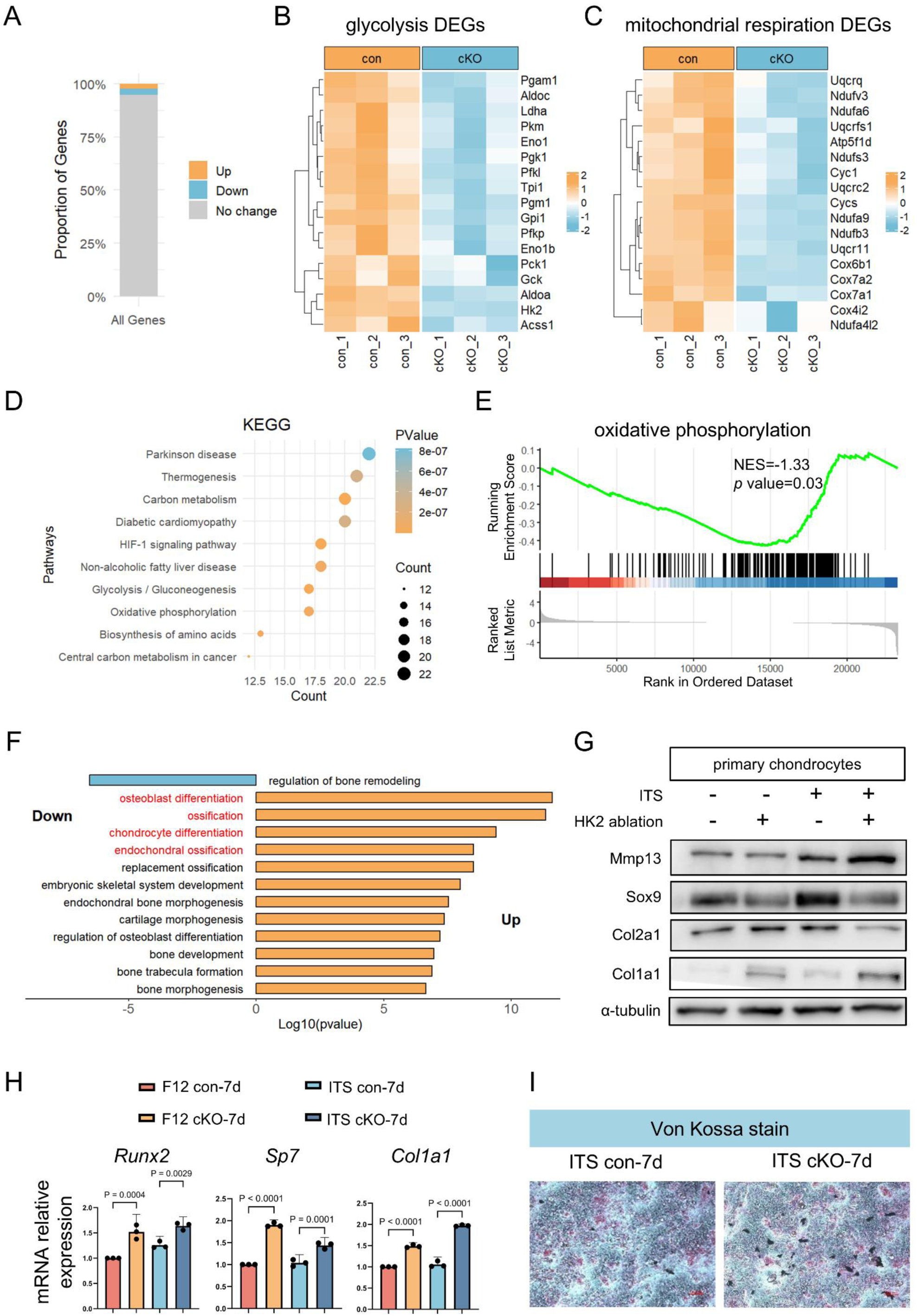
HK2 ablation in chondrocytes suppresses energy generation and promotes calcification. (A) Proportion of DEGs identified from bulk RNA-seq. (B) Heatmap of glycolysis-related gene expression. Color scale represents expression levels. (C) Heatmap of mitochondrial function-related gene expression. Color scale represents expression levels. (D) Bubble plot showing the top 10 downregulated KEGG pathways. (E) GSEA of the oxidative phosphorylation pathway. (F) Bar plot showing top significantly enriched bone and cartilage-related GO terms. (G) Immunoblotting analysis of Mmp13, Sox9, Col2a1 and Col1a1 protein levels in HK2-deficient chondrocytes compared to control cells. Cells were incubated with ITS for 7 days. α-tubulin was used as an internal control. (H) Relative mRNA expression levels of *Runx2*, *Sp7* and *Col1a1* in cells described in (g). Data are presented as mean ±SEM. *P < 0.05, **P < 0.01 indicates significant differences. (I) Representative images of Von Kossa staining in HK2-deficient chondrocytes and control cells. Chondrocytes were induced by ITS for 7 days.

Interestingly, Gene Ontology (GO) analysis revealed that, despite pathways related to energy generation and nucleotide metabolism being suppressed (Fig. S1B, C), genes involved in osteoblast differentiation, ossification, chondrocyte differentiation, endochondral ossification, and replacement ossification were significantly upregulated (P < 0.0001) (Fig. 2F), suggesting a probable promotion in cartilage degradation and calcification. Consistent with these findings, the relative mRNA levels of *Col1a1* (P <0.0001), *Runx2* (P = 0.0028) and *Sp7* (P = 0.0027), key regulators of cartilage calcification, were significantly increased in HK2-deficient chondrocytes (Fig. 2G). Immunoblotting analysis further confirmed a reduction in protein levels of Sox9 and Col2a1 but an elevation in Runx2 and Col1a1, regardless of induction conditions (Fig. 2H). Given that Runx2 is a critical transcription regulator of chondrocyte hypertrophy [27], and hypertrophic chondrocytes are capable of producing calcium-containing crystals involving calcification [1, 28], we next performed Von kossa staining to assess chondral calcification. As expected, mineralized nodule formation was significantly increased in HK2-deficient chondrocytes after 7 days of culture compared with intra-group controls (Fig. 2I). These findings suggest that HK2 ablation suppress chondrogenic differentiation and leads to promotion in cartilage calcification.

### A glance of HK2 expression in chondrocytes across conditions

Cartilage calcification is a hallmark of OA[28]. To identify a pathological role of HK2, we analyzed population heterogeneity and reconstructed the reprogramming trajectory of human knee cartilage cells using an open-access single-cell transcriptomic dataset (GSE255460) [29]. Cells were ordered in pseudotime using Monocle 2, an algorithm for the lineage reconstruction of biological processes based on transcriptional similarity [30]. Notably, cell clusters from weight-bearing cartilage in OA samples (WB, PC=18, resolution =0.8) differed markedly from those in non-weight-bearing OA cartilage (NWB, PC=14, resolution=0.6) or non-OA controls (PC=15, resolution =0.8) (Fig. 3A). In non-OA cartilages, the differentiation trajectory bifurcated into two terminal fates: hypertrophic/fibrocartilage chondrocytes and proliferative chondrocytes (Fig. 3B). And in NWB OA cartilage, the differentiation trajectory bifurcated into hypertrophic/fibrocartilage chondrocytes and reparative chondrocytes (Fig. 3C). However, WB OA cartilage showed divergence into hypertrophic/fibrocartilage chondrocytes and inflammatory chondrocytes (Fig. 3D). These findings indicate that the chondrocyte fate undergoes significant alterations under diverse situations. Notably, along the pseudotime trajectory into hypertrophic/fibrosis fate across conditions, HK2 expression in healthy cartilage was high at early differentiation stages, but declined sharply at intermediate stages, and remained low at terminal stages (Fig. 3E), which consistent with the results in Figure 1A. However, in OA cartilages, HK2 expression was relatively low during early differentiation stages but increased toward the terminal stages (Fig. 3F, G) and remained gradually elevated throughout the trajectory in the fate which is responsible for the differentiation of inflammatory chondrocytes (Fig. 3H). These findings suggest that HK2 may participate in determining chondrogenic fate, in the context of alterations in mechanical stress or inflammation.

**Fig. 3.**
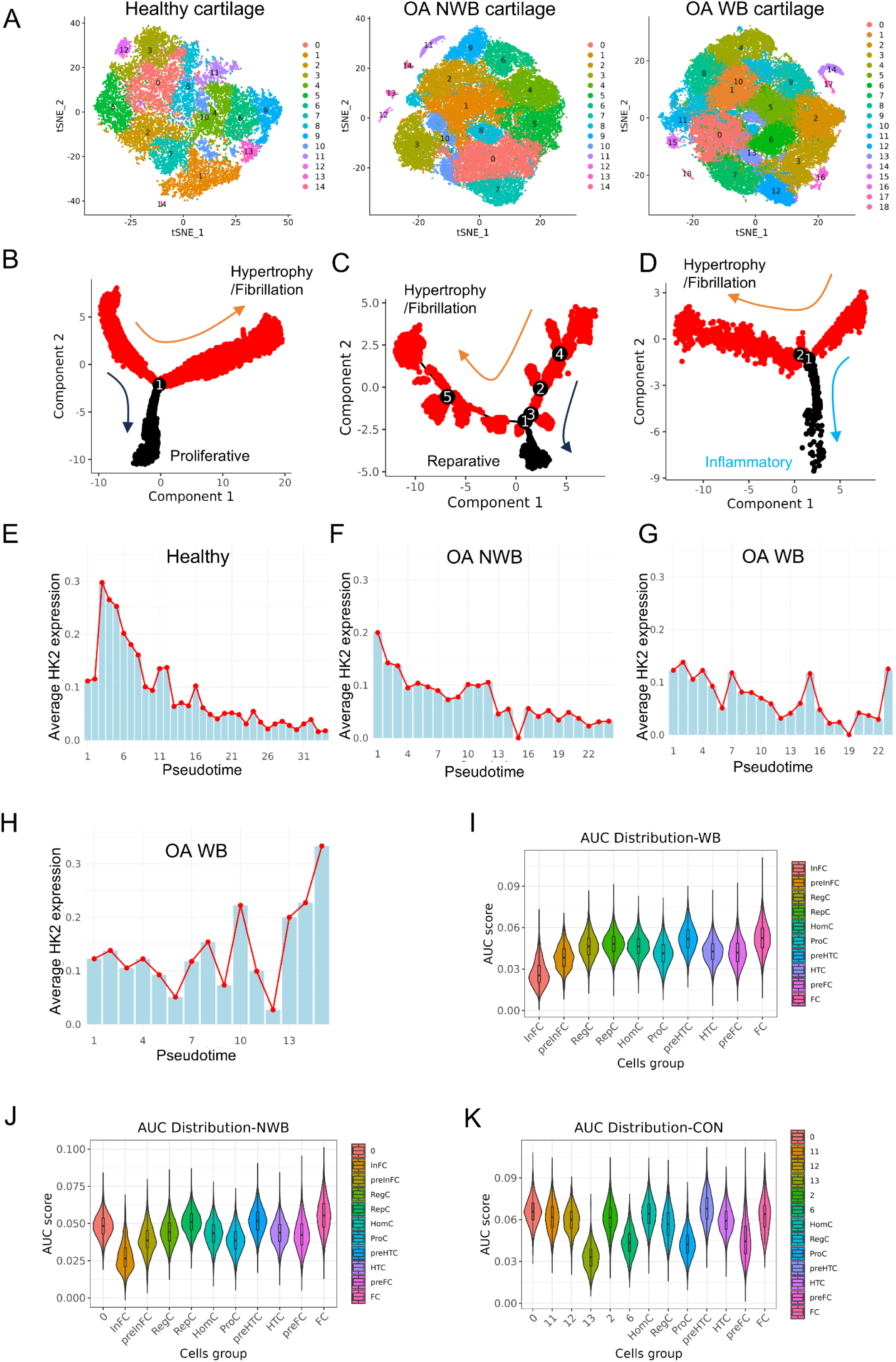
A glance of HK2 expression in chondrocytes across conditions. (A) t-SNE plot showing transcriptional profiles of chondrocytes from healthy (n = 19,225), OA non-weight-bearing (OA NWB, n = 60,616), and OA weight-bearing (OA WB, n = 67,033) cartilages. (B-D) Pseudotime trajectory of chondrocytes inferred by Monocle2 in (B) healthy, (C) OA NWB, and (D) OA WB cartilages. (E-G) HK2 expression dynamics along the hypertrophy/fibrillation branch in (E) healthy, (F) OA NWB, and (G) OA WB cartilage. (H) HK2 expression dynamics along the inflammatory branch in OA WB cartilages. (I-K) AUCell analysis of upregulated DEGs from bulk RNA-seq overlaid on scRNA-seq data. (i) OA WB, (j) OA NWB, and (k) healthy cartilages.

Importantly, by mapping the upregulated differential expression genes (DEGs) from bulk RNA-seq onto the scRNA-seq datasets in GSE255460, we identified that majority of the DEGs are expressed in clusters of pre-hypertrophic and fibrotic chondrocytes from OA (Fig. 3I, J) and healthy cartilages (Fig. 3K), providing evidence to support our hypothesis on a probable contribution of HK2 in chondrocyte fate.

### 3-BrPA and HK2 ablation both aggravate OA progression in mice

To evaluate the effects of chondrocyte HK2 in pathological conditions, we first employed a DMM-induced OA model in mice and administered with 3-BrPA. We determined that HK2 expression was significantly reduced in articular cartilages of DMM-operated mice compared to sham controls (Fig. 4A). Mice were administered intraperitoneal injections of 3-BrPA (5 mg/kg) twice weekly for six weeks post-surgery (Fig. 4B). Safranin O staining and OARSI scoring revealed aggravated cartilage destruction in 3-BrPA-treated OA mice (P = 0.0321) (Fig. 4C, D). Consistently, 3-BrPA treatment resulted in a significant decrease in Col2a1 expression (P = 0.0001) (Fig. 4E) and an increase in Mmp13 expression (P = 0.0001) (Fig. 4F) in articular cartilage under both sham and OA conditions. Additionally, TRAP staining showed enhanced chondroclast and osteoclast activity following 3-BrPA treatment in both groups (P = 0.0002) (Fig. 4G, H), indicating a potential outcome in subchondral bone destruction.

**Fig. 4.**
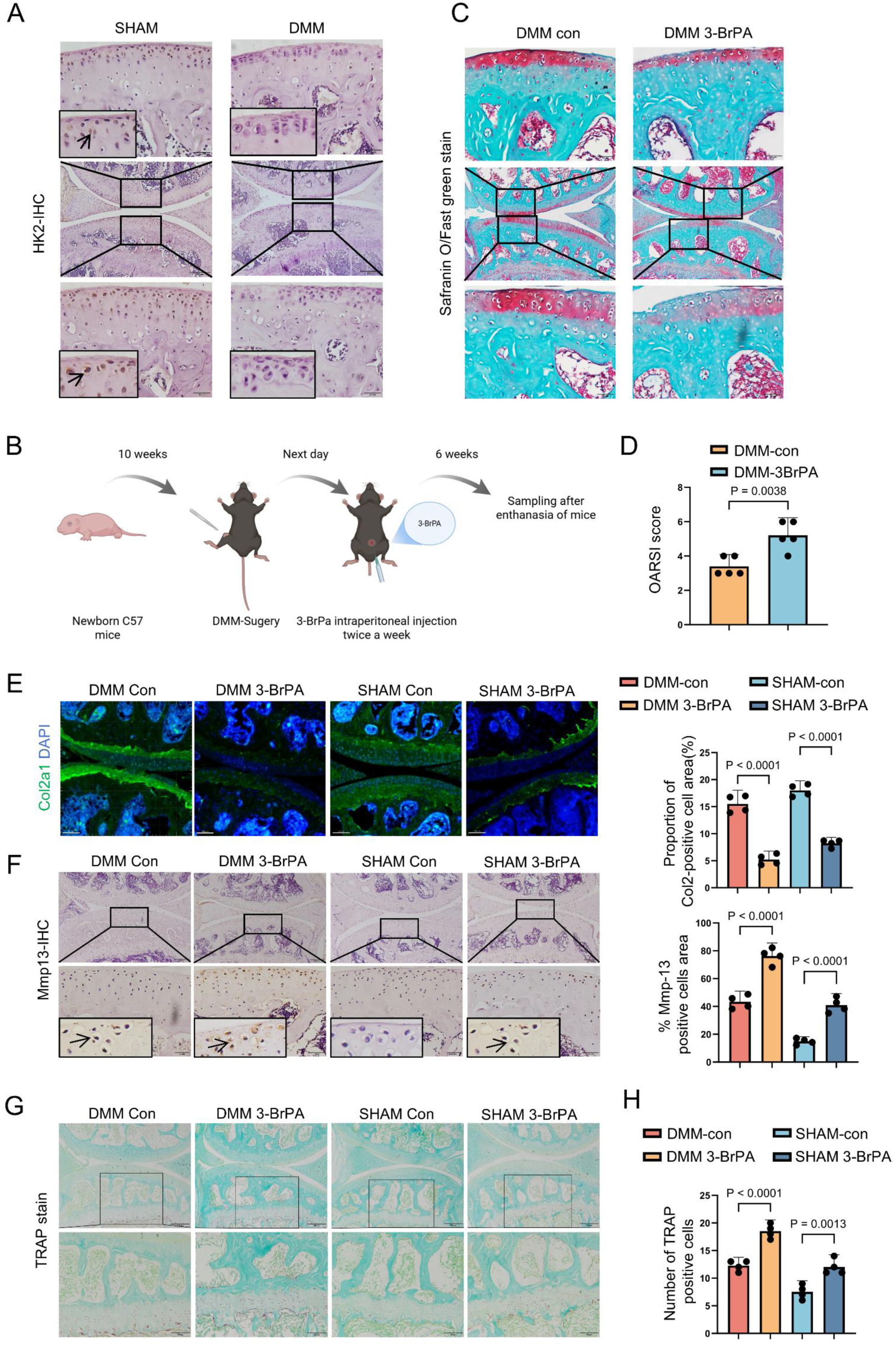
3-BrPA accelerates OA progression in mice. (A) Immunohistochemistry (IHC) staining of HK2 in articular cartilage following DMM-surgery. Scale bar = 100 μm, 20 μm. (B) Schematic diagram showing intraperitoneal injection of 3-BrPA (5 mg/kg) in DMM-surgery mice. (C) Representative images of Safranin O/Fast Green staining of articular cartilage from OA mice administered with or without 3-BrPA, scale bar = 100 μm, 20 μm. (D) OARSI scores comparing cartilage damage between 3-BrPA-treated mice and normal saline-treated controls (n = 5). (E) Immunofluorescence (IF) staining analysis of Col2a1 protein levels in articular cartilage from DMM mice administered with or without 3-BrPA (n = 4). Scale bar = 100 μm. (F) IHC staining analysis of Mmp13 levels in articular cartilage from DMM mice administered with or without 3-BrPA (n = 4). Scale bar = 200 μm, 20 μm. (G) TRAP staining analysis of chondroclast and osteoclast activity affected by 3-BrPA administration (n = 4). Scale bar = 100 μm, 50 μm. All data are presented as mean ± SEM. *P < 0.05, **P < 0.01 indicates significant differences.

Intriguingly, the whole-mount skeletal staining of 3-week-old mice (Fig. 5A), growth plate thickness measurement in 10-week-age mice (P = 0.1219) (Fig. 5B) and longitudinal growth monitoring (P =0.5325) (Fig. 5C, D) revealed no significant differences in skeletal development between cKO mice and their littermate controls. However, although previous studies have reported that elevated HK2 level correlating with disease severity in RA [8], mice with HK2-deficeint chondrocytes in the DMM-indued OA conditions exhibited exacerbated cartilage loss, as evidenced by safranin O staining (Fig. 6A) and increased OARSI scores (P = 0.0065) (Fig. 6B). Moreover, nociceptive sensitivity was significantly increased in cKO mice (P = 0.0072) (Fig. 6C), together suggesting that HK2 deletion in chondrocytes accelerates OA progression. Consistent with these findings, immunofluorescence analysis showed reduced Col2a1 expression in articular chondrocytes of cKO mice compared to controls under OA conditions (P = 0.0037) (Fig. 6D), while immunohistochemistry revealed elevated Mmp13 expression (P = 0.0011) (Fig. 6E), indicating a marked loss of hyaline cartilage matrix and enhanced degradation. Additionally, chondroclast and osteoclast activity were increased in cKO mice without surgery and further intensified post-DMM, parallel to the effects observed in 3-BrPA-treated OA mice (P = 0.0001) (Fig. 6F), indicating a promoting effect on subchondral bone destruction due to the absence of HK2 is further amplified in the context of OA.

**Fig. 5.**
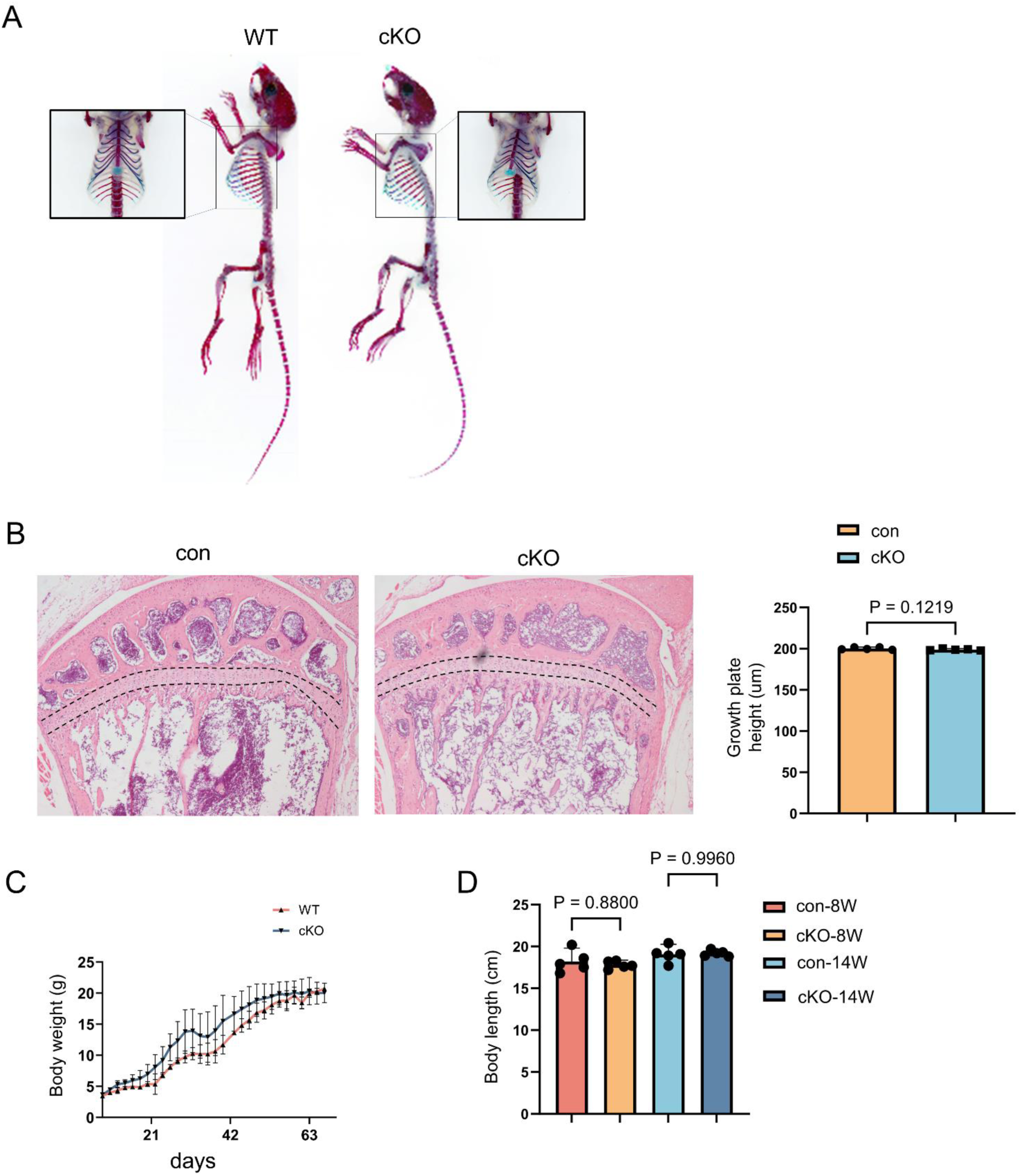
HK2-deficient chondrocytes do not affect skeletal development in mice. (A) Whole-mount skeletal staining of male 3-week-old cKO mice and their littermate controls. (B) Representative H&E staining showing the growth plate thickness, n.s.=non-significant. (C) Changes in body weight over time, each point represents the average body weight for each group, and the trend is shown. (D) Comparison of 8-and 14-week-old male cKO mice and their littermate controls. n.s.=non-significant.

**Fig. 6.**
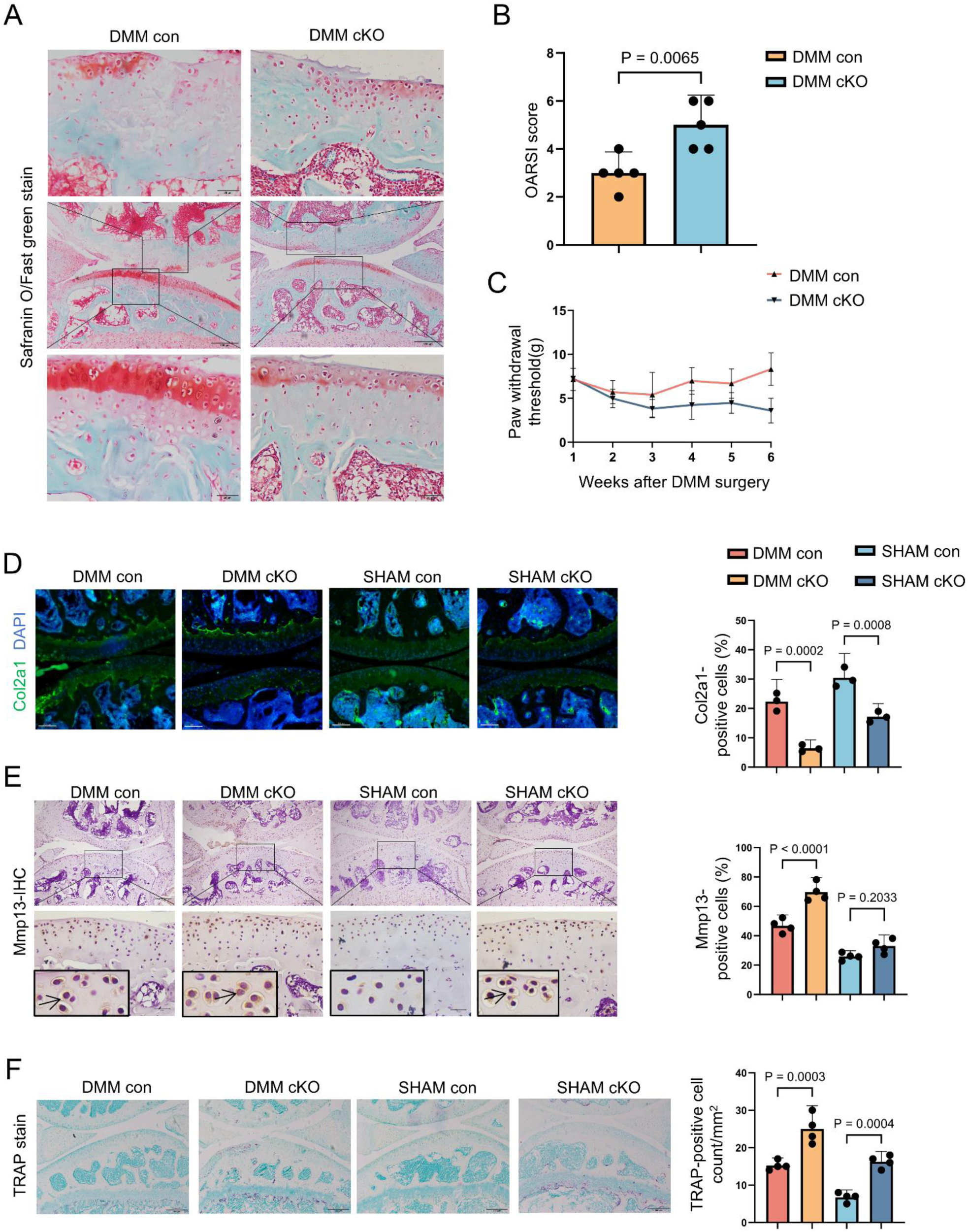
HK2-deficient chondrocytes promote OA development. (A) Representative images of Safranin O/Fast green staining of articular cartilage from DMM-cKO mice and their littermate controls (n = 5). Scale bar = 100 μm, 20 μm. (B) OARSI scoring of articular cartilage destruction in (A). (C) Pain sensitivity evaluation in DMM-cKO mice and their littermate controls (n = 5). (D) IF staining analysis of Col2a1 expression levels in articular cartilage from DMM-cKO and their littermate controls (n = 3). Scale bar = 100 μm. (E) IHC staining analysis of Mmp13 levels in articular cartilage from DMM-cKO and their littermate controls (n = 4). Scale bar = 200 μm, 20 μm. (F) TRAP staining analysis of chondroclast and osteoclast activity affected by HK2-deficeint chondrocytes in DMM-OA mice (n = 4). Scale bar = 100 μm. All data are presented as mean ± SEM. *P < 0.05, **P < 0.01 indicates significant differences.

Collectively, our findings suggest that HK2 expression is essential for chondrocytes to maintain homeostasis particularly under pathological conditions. Unlike RA, targeting HK2 as a therapeutic strategy for OA may pose significant risks. Under metabolic stress, chondrocytes may deprioritize biosynthetic and genomic activities associated with their chondrogenic identity in favor of bone-like differentiation and calcification. Further studies are warranted to clarify the molecular mechanisms through which HK2 regulates these processes.

## Discussion

Metabolic enzymes play a fundamental role across diverse cell types by regulating energy metabolism to support cellular functions and maintain tissue and organ integrity. In OA, chondrocytes exhibit reduced mitochondrial respiration and increased glycolysis, a metabolic shift thought to contribute significantly to OA progression [4, 5, 31]. Although emerging evidence suggests that HK2 may be a promising therapeutic target in rheumatoid arthritis (RA) [32], our findings indicate that targeting HK2 in chondrocytes impairs cartilage function and exacerbates OA pathology.

Bone and cartilage calcification are physiologically regulated processes [33]. However, cartilage calcification is also observed in pre-osteoarthritic joints [34] and is considered a hallmark of degenerative joint disease [28, 35]. Despite its clinical significance, the regulatory mechanisms initiating pathological calcification remain largely elusive. In OA, articular chondrocytes display features similar to those in the growth plate, including chondral calcification [36, 37], suggesting a possible shared mechanism between OA-related cartilage degeneration and endochondral ossification [38]. Interestingly, in our study, HK2 deficiency in chondrocytes accelerated OA progression without affecting growth plate development, despite HK2 ablation impairing chondrogenesis ex vivo. These distinct outcomes imply that HK2 inhibition may contribute to the initiation of pathological calcification in articular cartilage.

HK2 functions not only as a rate-limiting enzyme in glycolysis but also interacts with mitochondria via its N-terminal helix [39], thereby enhancing mitochondrial glucose utilization. Experimental studies have demonstrated essential roles for mitochondrial HK2 in various pathological conditions. In the brain, HK2 plays dual roles in regulating microglial function by modulating both glycolytic flux and mitochondrial activity [40]. Mitochondrial dissociation of HK2 can trigger mitophagy and confer cardioprotection during ischemic injury [41]. Beyond its metabolic role, HK2 has also been found to localize to the nucleus, where it helps maintain stemness in hematopoietic and leukemic stem cells [22]. These diverse subcellular localizations underscore the context-dependent and multifaceted functions of HK2 in both metabolic and non-metabolic pathways.

In conclusion, our findings reveal a crucial role for the glycolytic enzyme HK2 in chondrocyte differentiation. Considering that pathological cartilage calcification is a prominent feature of joint injury, and that aging, mechanical stress, inflammation, metabolic disorders, and hormonal factors can disrupt cartilage homeostasis [42], HK2 may serve as a critical integrator of cartilage integrity, particularly under stress or pathological conditions. Elucidating how HK2 coordinates these distinct cellular processes may offer valuable insights into OA pathogenesis and identify potential therapeutic avenues.

## Supporting information

Primers

## List of abbreviations

DEGs: differential expression genes
DMM: destabilization of the medial meniscus
GSEA: gene set enrichment analysis
GO: Gene Ontology
HK2: hexokinase 2
KO: knockout
KEGG: Kyoto Encyclopedia of Genes and Genomes
NWB: non weight bearing OA cartilage
OA: osteoarthritis
OARSI: Osteoarthritis Research Society International
OXPHOS: oxidative phosphorylation
RA: rheumatoid arthritis
scRNA-seq: single cell RNA sequencing
WB: weight bearing cartilage
3-BrPA: 3-bromopyruvate

## Declarations

### Ethics approval and consent to participate

All animal procedures were approved by the Southern Medical University Animal Care and Use Committee and conducted in accordance with institutional ethical guidelines.

### Availability of data and materials

Data supporting the findings of this study are available within the article. The single-cell RNA-seq dataset used in this study is publicly available in the NCBI’s Gene Expression Omnibus (GEO) under accession number GSE255460, originally published by Fan Y, Bian X, Sun S et al. (2024).

## Acknowledgements

All authors gratefully thank Prof. Xiao Yang (Academy of Military Medical Sciences, China) for kindly proving the Col2a1-cre mice line. All authors acknowledge the authors and contributors of the dataset [GSE255460], available from the GEO, for making their data publicly accessible.

## Authorship contributions

Haokun Xu: Investigation, Methodology, Data curation, Formal analysis. Xinyi Zhang: Bioinformatics analysis, Validation, Data curation, Formal analysis. Yijie Fu: Investigation. Sirui Yuan: Investigation. Daizhao Deng: Investigation. Ying Lin: Investigation. Ruijun Lai: Investigation. Kai Li: Methodology & Review. Tinghui Xiao: Methodology & Review. Song Xu: Review & Funding acquisition. Xiaochun Bai: Project administration. Yue Zhang: Writing – original draft, review & editing, Project administration, Funding acquisition, Conceptualization.

