## Supplementary material for "Inhibition of hexokinase 2 undermines cartilage health and accelerates osteoarthritis": Figure S1

### Supplemental Figure and Figure legend

Figure S1

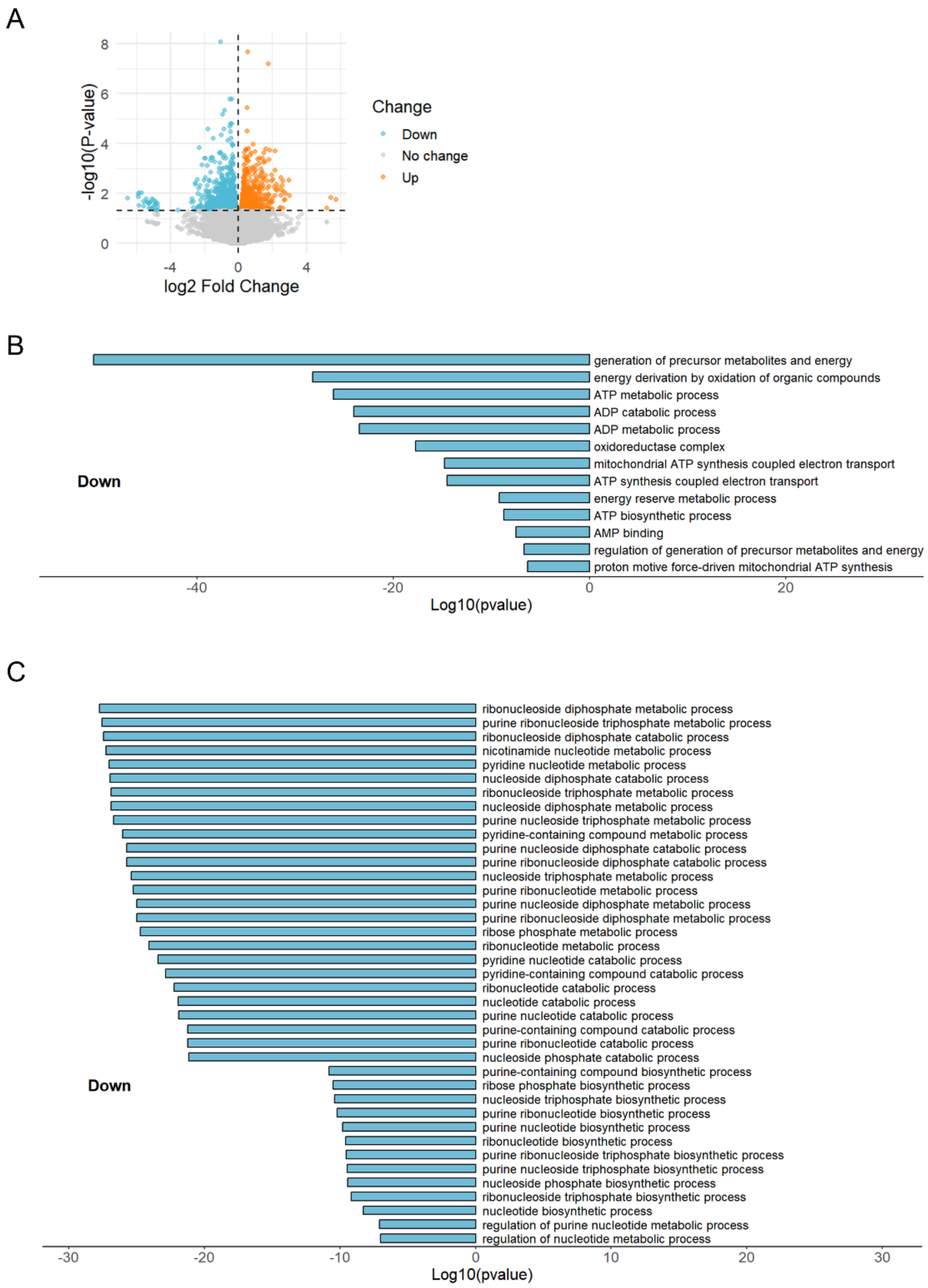


**Figure S1. Bulk RNA-seq analysis of HK2-deficient primary chondrocytes**

**A.** Volcano plot illustrates differentially expressed genes in HK2-deficient primary chondrocytes.

**B.** Bar plot showing the top significantly enriched GO terms related to energy generation.

**C.** Bar plot showing the top significantly enriched GO terms associated with nucleotide metabolism.
